# Chronic administration of a positive allosteric modulator at the α5-GABAA receptor reverses age-related dendritic shrinkage

**DOI:** 10.1101/2020.02.28.964742

**Authors:** Thomas D. Prevot, Akiko Sumitomo, Toshifumi Tomoda, Daniel E. Knutson, Guanguan Li, James M. Cook, Etienne Sibille

## Abstract

Over the last 15 years, worldwide life expectancy increased by 5 years jumping from 66 years to 71 years. With progress in science, medicine, and care we tend to live longer. Such extended life expectancy is still associated with age-related changes, including in the brain. The aging brain goes through various changes that can be called morphomolecular senescence. Overall, the brain volume changes, neuronal activity is modified and plasticity of the cells diminishes, sometimes leading to neuronal atrophy and death. Altogether, these changes contribute to the emergence of cognitive decline that still does not have an efficient treatment available. Many studies in the context of cognitive decline focused on pathological aging, targeting β-amyloid in Alzheimer’s disease, for example. However, β-amyloid plaques are also present in healthy adults and treatments targeting plaques have failed to improve cognitive functions. In order to improve the quality of life of aging population, it is crucial to focus on the development of novel therapies targeting different systems altered during aging, such as the GABAergic system. In previous studies, it has been shown that positive allosteric modulators (PAM) acting at the α5-containing GABA-A receptors improve cognitive performances, and that these α5-GABA-A receptors are implicated in dendritic growth of pyramidal neurons. Here, we hypothesized that targeting the α5-GABA-A receptors could contribute to the reduction of cognitive decline, directly through activity of the receptors, and indirectly by increasing neuronal morphology. Using primary neuronal culture and chronic treatment in mice, we demonstrated that an α5-PAM increased dendritic length, spine count and spine density in brain regions involved in cognitive processes (prefrontal cortex and hippocampus). We also confirmed the procognitive efficacy of the α5-PAM and showed that the washout period diminishes the precognitive effects without altering the effect on neuronal morphology. Future studies will be needed to investigate what downstream mechanisms responsible for the neurotrophic effect of the α5-PAM.

## Introduction

Thinking about cognitive decline in aging, the first things that come in mind are Alzheimer’s disease and related dementias, but human and animal suffer from cognitive decline during normal aging without necessarily developing neurodegenerative diseases. With progress in medicine and health care, human population tends to live longer. Since 2018, persons aged 65 or above have outnumbered children under 5 years of age globally (1, 2). By 2050, it is estimated that 16% of the global population will be over 65 years of age, while in 2019 only 9% of the population is over 65 years of age (2, 3). Overall, such extended lifespan is accompanied by normal cognitive decline and brain changes.

Normal aging has been studied for decades and it is now acknowledged that neurocognitive, structural, functional and cellular changes are hallmarks of aging (4). Cognitive changes with aging are not necessarily associated with poorer functions. For example, vocabulary use or general knowledge and expert skills remain intact or can even improve with aging (5, 6). In the other hand, other functions such as problem solving (7), memory (8, 9), processing speed (10) and psychomotor abilities (11) (also known as executive functions) suffer gradual impairment with increasing age. More specifically, episodic memory (12) (memory of personal experienced events) and working memory (13) (temporary maintenance, storage and update of information while performing a task) are altered during aging (8).

Cognitive decline was linked with brain changes at multiple levels that can be defined as “morphomolecuclar senescence”(14). Aging causes a grey matter volume decline, particularly in the frontal areas of the cortex (anterior cingulate cortex: (15)). Subcortical regions including the hippocampus, brain region critically involved in cognitive processes, also suffer from volume atrophy (16). Such decrease in grey matter in frontal areas and in the hippocampus has been suggested to be caused by neuronal shrinkage and death (17, 18), altogether contributing to cognitive decline. Pyramidal neurons, excitatory neurons in cortical layers, are particularly vulnerable during aging (19-21), and exhibit reduced spine density and dendritic shrinkage during normal aging. This neuronal atrophy has been showed across species, from human post mortem studies (22) to rodents (23), including monkeys (24). Multiple factors contribute to neuronal shrinkage during normal aging, including, neuroinflammation (25) accumulation of β-amyloid (26–29) or dysregulation of neuronal excitation-inhibition (21).

Neuroinflammation and β-amyloid load have been extensively studied, especially in the context of Alzheimer’s disease (30–32). However, more and more studies are showing that amyloid load is a normal process during aging (29), and the relationship between amyloid load and cognitive decline remains uncertain in healthy elderly (33, 34). Multiple pharmaceutical companies tried to develop treatment by targeting β-amyloid plaques(35), with very limited success to date (36). To increase the changes of success at countering cognitive decline, scientists have to explore other paths than β-amyloid.

Neuronal activity and the excitation/inhibition balance is also impaired during normal and pathological aging. Some studies have shown conflicting results sometimes suggesting a decreased excitability of neurons toward a hyper-inhibition (37, 38), and sometimes showing overall decrease in both excitation and inhibition (39–41). GABAergic neurons represents the major inhibitory players to control excitability of excitatory cells. Using magnetic resonance spectroscopy, GABAergic activity has been shown to be downregulated during normal and pathological aging, and such deficit in GABA was associated with cognitive decline (42). GABA signal through two main types of receptor, known as GABA-A and GABA-B receptors. The GABA-A receptors are ion-channel receptors that hyperpolarize the post-synaptic cell by chloride influx (43). The GABA-A receptors are pentameric, composed of a combination of 5 subunits between α, β, γ, δ, ε, θ, π and ρ (44). The different subunits of the GABA-A receptors show variable dysregulation with aging, depending on the brain regions (45). Some studies showed downregulation of α_1_, β_1_, β_2_, γ_1_ and γ_2_ subunits in the rat auditory cortex (46) and others showed that α_3_ and α_5_ subunits were downregulated in the rat motor cortex (47). Other studies also showed an upregulation of the α_1_-subunit in the hippocampus of old rats (48), while a decrease of the same subunit was observed in monkeys (49). Despite variability in the changes of the GABA-A receptors associated with age, all studies agree that decrease in the number of GABA-A receptors can result in behavioral changes (50), including cognitive deficits (51). The α_5_-subunit has a particular distribution in the brain, as it is located exclusively in brain regions highly involved in cognitive processes, such as the hippocampus and the prefrontal cortex (52). Preclinical and clinical data showed that the α_5_-subunit is downregulated in aging (53–55), and a recent study showed that synaptic localization of α_5_-containing GABA-A receptors contributes to dendritic outgrowth and spine maturation (56). Altogether, this data suggests a critical role of this particular subunit in the regulation of cognitive function mediated by the GABAergic system, directly through reversal of decreased activity of the receptor during aging, or indirectly through dendritic growth.

Benzodiazepine, such as diazepam, bind at the interface between the α_1,2,3,5_-subunit and the γ-subunit (57), act as a positive allosteric modulator (PAM) and are the most commonly prescribed medication for the treatment of anxiety, widely prescribed to elderly despite apparent issues and complications linked with age (58). The wide range of activity at different subunits is responsible for considerable side effects that can be potentiated by the GABAergic dysregulation occurring during normal or pathological aging (48). Novel developments have tried to reduce the side effects of novel benzodiazepines, or derivatives, by increasing selectivity to the α_5_-subunit and reducing selectivity to the α_1_-subunit (59). Previous data from our group showed that the α_5_-positive allosteric modulator (α_5_-PAM) GL-II-73 improves working memory in mouse model of chronic stress and aging (59). Other group demonstrated improved cognitive functions in old rats using different α_5_-PAMs, namely Compound 44 and Compound 6 (60). Knowing the distribution of the α_5_-subunit in the brain, its downregulation during aging, its potential role in dendritic growth, and the known procognitive effect of GL-II-73 in old mice, we hypothesized that facilitating the action of GABA at the α_5_-GABA receptors with GL-II-73 will improve cognitive functions in old mice, directly through activity at the receptor, and indirectly through reversal of dendritic shrinkage due to normal aging. To validate our hypothesis, we used primary neuronal culture from mouse hippocampus to test the effect of GL-II-73 *in vitro*, at promoting dendritic growth. We then tested the effect of GL-II-73 *in vivo*, in old mice, to demonstrate if chronic treatment can improve cognitive deficits and reverse dendritic shrinkage, and if such effects are maintained when the treatment stops.

## Material and Methods

### Animals

For the cell culture experiment, heterozygote CamKII-Cre male mice (Jackson Laboratories, Cat#5359) were crossed with Rosa26-flox-stop-GFP female mice to express the green fluorescent protein (GFP) exclusively in pyramidal neuronal. The pregnancy of the females was timed, in order to collect embryos at E14 for primary cortical neuronal culture. Pups were quickly decapitated, and the prefrontal cortex was harvested for primary neuronal culture. For the behavioral experiment, two separate cohorts of 30 male C57BL6 mice were purchased as retired breeders from Charles River Laboratories at the age of 9-10 months, and kept in the animal facility until they reach the age of 22 months. Two cohorts of young male C57BL6 mice (N1=11; N2=8) were purchased separately to include a group of young mice for each cohort of old mice. Seven retired breeders from the first cohort, and 12 from the second cohort died of normal aging before the initiation of the behavioral tests.

### Primary cortical neurons in culture and morphometric analysis

To visualize dendritic and spine morphology of major output neurons in culture, we prepared primary cortical neurons (61) and maintained the culture for up to 25 days by replacing half the media (Neurobasal media supplemented with B-27 [GibcoBRL]) every other day. To evaluate dendritic branching complexity, ImageJ with Sholl analysis plug-in (http://fiji.sc/Sholl_Analysis) is used to score the number of intersections of dendritic arbors with a series of concentric spherical shells drawn from the soma with periodic distance. Numbers of spines along all apical dendritic branches (except for the main shaft of apical dendrite) per cell are scored by ImageJ with bio-formats plug-in, and the average spine numbers per 100 μm length are calculated.

### Drug preparation

The positive allosteric modulator (PAM) at the α_5_-containing GABA-A receptor (α_5_-PAM) GL-II-73 was synthesized in collaboration with Dr. Cook’s group (University of Wisconsin-Milwaukee). For the neuronal culture, the α5-PAM GL-II-73 was infused in the media at the concentration of 1µM, in 0.01% DMSO. The vehicle solution was only 0.01% DMSO, mixed in the media. The drug was left incubating in the culture for 24 hrs. For the behavioral studies, GL-II-73 was administered through the drinking water at a dose of 30mg/kg/day, for 30 days, based on previous studies from our group (59).

### Open Field

Mice were habituated to the room lit at 75lux for 30 minutes prior to testing. The apparatus was a grey PVC arena (43×43cm) with walls 43cm high. A digital camera was mounted to record the animals’ activity in the arena for 10 minutes. Post-acquisition videos were analyzed using the software ANYmaze (Stoelting). Distance travelled in meters (m) was measure as a proxy for locomotor activity.

### Y-Maze Alternation Task

The apparatus is made of black PVC, shaped like a Y, with a sliding door at the entry of each arm. Mice were habituated to the maze, by letting them explore the maze freely for 10 minutes per day, for 2 days. On the third day, mice were trained to alternate in the maze. Each animal was placed in the starting box for 30 seconds, before opening of the door. Once the door was opened, the animal could decide to visit the right or left arm of the maze. Once chosen, the animal was confined into that arm for 30 seconds. Then, the animal was gently transferred back to the starting box, with 30 seconds inter-trial-interval (ITI), prior to the next trial, identical to the previous one. If the animal did not alternate over 3 consecutive trials, the animal was then forced to alternate in order to prevent any reinforcement of one arm. This sequence was followed for 7 trials. Finally, on the fourth day, animals are subjected to the same sequence of testing but with a 60 sec ITI. Also, an 8^th^ trial with a 5sec ITI is implemented to assess potential loss of motivation. In the event of a lack of alternation at the 8^th^ trial, mice were removed from the statistical analysis.

### Brain collection and staining

Twenty-four (24) hours after completion of the last behavioral test, mice were euthanized using cervical dislocation, and their brains were harvested for downstream analysis. Four brains per group were used for the Golgi-Cox staining. The remaining brains were kept as backups, or for follow-up analyses. The brain extraction and initial immersion of Golgi staining procedure were conducted in CAMH, with Golgi-Cox staining solutions provided by NeuroDigiTech LLC. The brain samples were then shipped to NeuroDigiTech for sectioning and blinded morphological analysis in layers II/III of the prefontal cortex (PFC) and CA1 region of the hippocampus (CA1).

### Selection of the regions of interest and samplings

Brains were sectioned on a cryostat, and slices were mounted on glass slides. The slides included serial coronal sections that covered the anterior-to-posterior axis of the brain. The sampling of the regions of interest (ROIs) included the basal and apical dendrites of pyramidal cells in Layers II/III of PFC and the CA1 of the hippocampus. The ROIs were then chosen and analyzed using the stereology-based software NeuroLucida v10 (Microbrightfield, VT), installed on a Dell PC workstation that controlled Zeiss Axioplan 2 image microscope with Optronics MicroFire CCD camera (1600 x 1200) digital camera, motorized X, Y, and Z-focus for high-resolution image acquisition and digital quantitation. The sampling process was conducted as follows: the investigators first previewed the entire rostro-caudal axis of ROIs, under low-mag Zeiss objectives (10x and 20x), compared and located those with the least truncations of distal dendrites as possible under high-mag Zeiss objectives (40x and 63x), and then used a Zeiss 100x objective with immersion oil to perform 3D dendritic reconstruction, followed by counting of the spines throughout the entire dendritic trees. The criteria for selecting candidate neurons for analysis (n=6 per animal) were based on visualization of a completely filled soma with no overlap of neighboring soma and completely filled dendrites, the tapering of most distal dendrites; the visualization of the complete 3-D profile of dendritic trees using the 3-D display of imaging software. Neurons with incomplete impregnation and/or neurons with truncations due to the plane of sectioning were not collected. Moreover, cells with dendrites labeled retrogradely by impregnation in the surrounding neuropil were excluded (derived from (62)).

For spine sampling, only spines orthogonal to the dendritic shaft were readily resolved and included in this analysis, whereas spines protruding above or beneath the dendritic shaft were not sampled. This principle was remained consistent throughout the course of analysis. After completion, the digital profile of neuron morphology was extrapolated and transported to a multi-panel computer workstation for the quantitative analysis, including the dendrograms, spine counts, and Sholl analyses.

### Experimental Design

In the first experiment, 11 young and 22 old mice were included. Among the 22 old mice, 8 received GL-II-73 in the drinking at the dose of 30mg/kg, while the other 14 received only water. This group design is due to the limited availability of the drug that did not allow more animals to be included in the treatment group. Similarly, in the second study, 8 young and 18 old mice were included. Among the old mice, 6 received water, 6 received GL-II-73 and 6 received GL-II-73 with a 1-week washout. All young mice from both behavioral experiments received water for the entire duration of the study. In each group, 4 mice were used to the Golgi staining analyses. Selection of the animals for this study was random, and ensured to be representative of the average performances of the group at large.

### Statistical analysis

In the cell culture experiment, the average number of intersection depending on the distance from the soma, and the treatment group was analyzed using 2-way ANOVA in repeated measure. Statistical differences were then characterized using *Scheffe’s test*. The average spine count per group was analyzed using a *t*-test (two-tailed). Behavioral performances were analyzed using 1-way ANOVA followed by *post hoc* analyses when significance was reached (*Scheffe’s test*). To quantify the dendritic length, spine count and spine density the samples, 6 cells per animal per brain region were used. The total of each feature was quantified, as well as the detail of each feature on the apical or basal segment of the dendrite. Repeated measure ANOVA taking into consideration each cell quantified for its dendritic length, spine count and spine density was performed with “group” as the independent variable. If a significant difference was found, Tukey’s multiple comparison test was performed to identify the differences between groups. Correlation analyses were also performed between quantification in the PFC and the CA1 using linear regression.

## Results

### Infusion of GL-II-73 in neuronal culture increases dendritic complexity

The α5-PAM GL-II-73 was infused in neuronal culture expressing GFP, and imaged after 24hr (Fig.1A). The number of intersection between dendrites and concentric circles 10µm apart from each other (Scholl analysis) were analyzed (Fig.1B). Repeated measure ANOVA on 10 neurons per group showed a significant effect of GL-II-73 (p<0.05). *Post hoc* analysis identified an increase in intersection in the neurons treated with GL-II-73 between 30 and 50 µm from the cell body. Analysis of the spine number also showed a significant increase in neurons treated with GL-II-73 compared to neurons treated with vehicle (p<0.001; Fig.1C).

**Figure 1.**
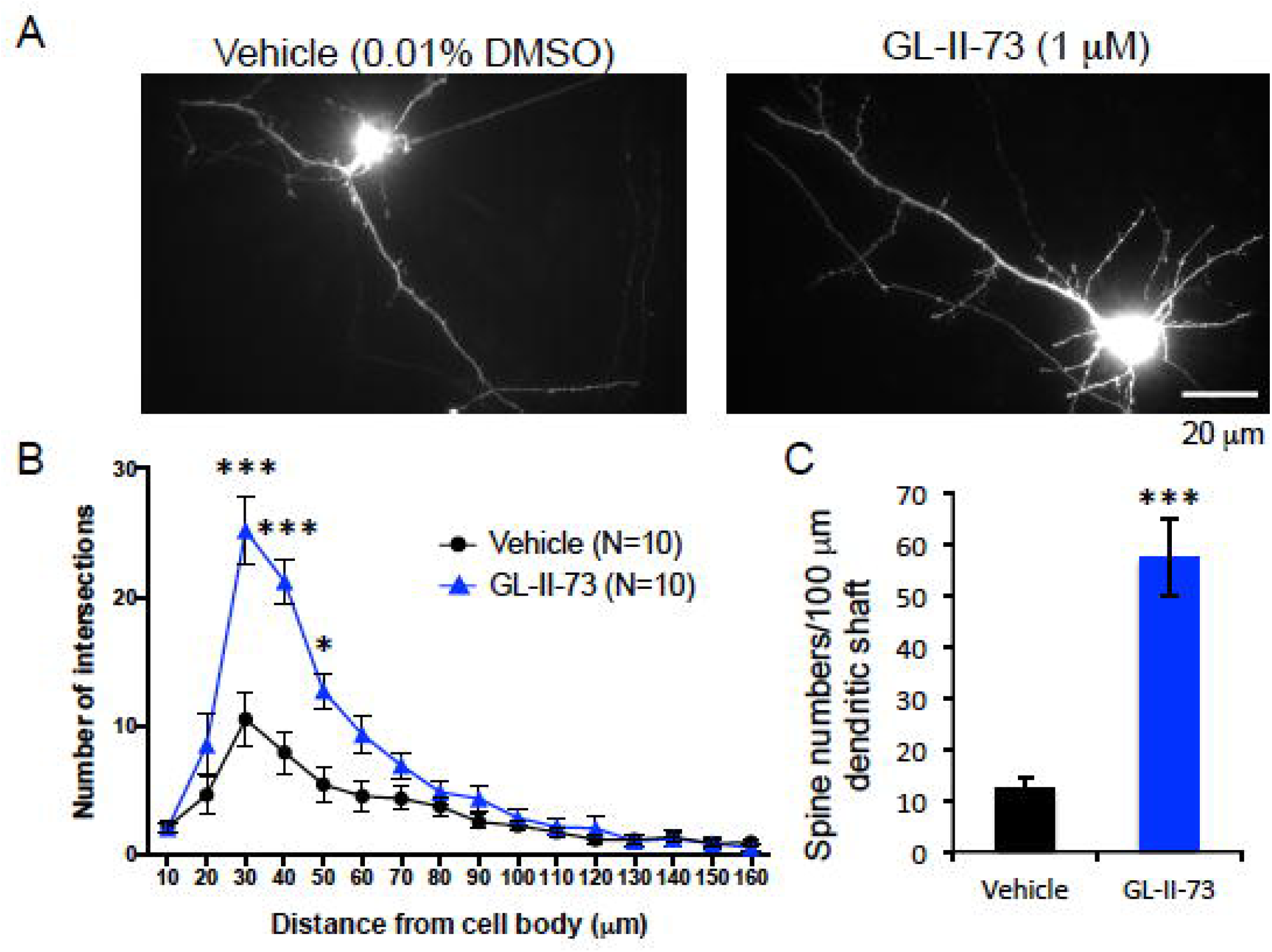
Effect of α_5_-PAM on dendrite morphology and spine numbers in primary cortical neurons in vitro. Cortical neurons were cultured from embryos of mice expressing the fluorescent protein GFP in pyramidal neurons (CamKII positive neurons). Neurons were cultured for 25 days and then incubated with Vehicle (0.01% DMSO in media) or GL-II-73 (1µM). (**A**) After 24hrs of incubation with GL-II-73, or vehicle, isolated neurons were imaged in order to analyze the morphology and the spine number. (**B**) Using the Sholl Analysis package of the software ImageJ, the number of intersections every 10µm from the soma was analyzed. A significant increase in the number of intersections was identified in the neurons incubated with GL-II-73 between 30 and 50 µm of distance from the soma. (**C**) Spine counts were also increased after incubation of GL-II-73. *p<0.05. ***p<0.001.

**Figure 2.**
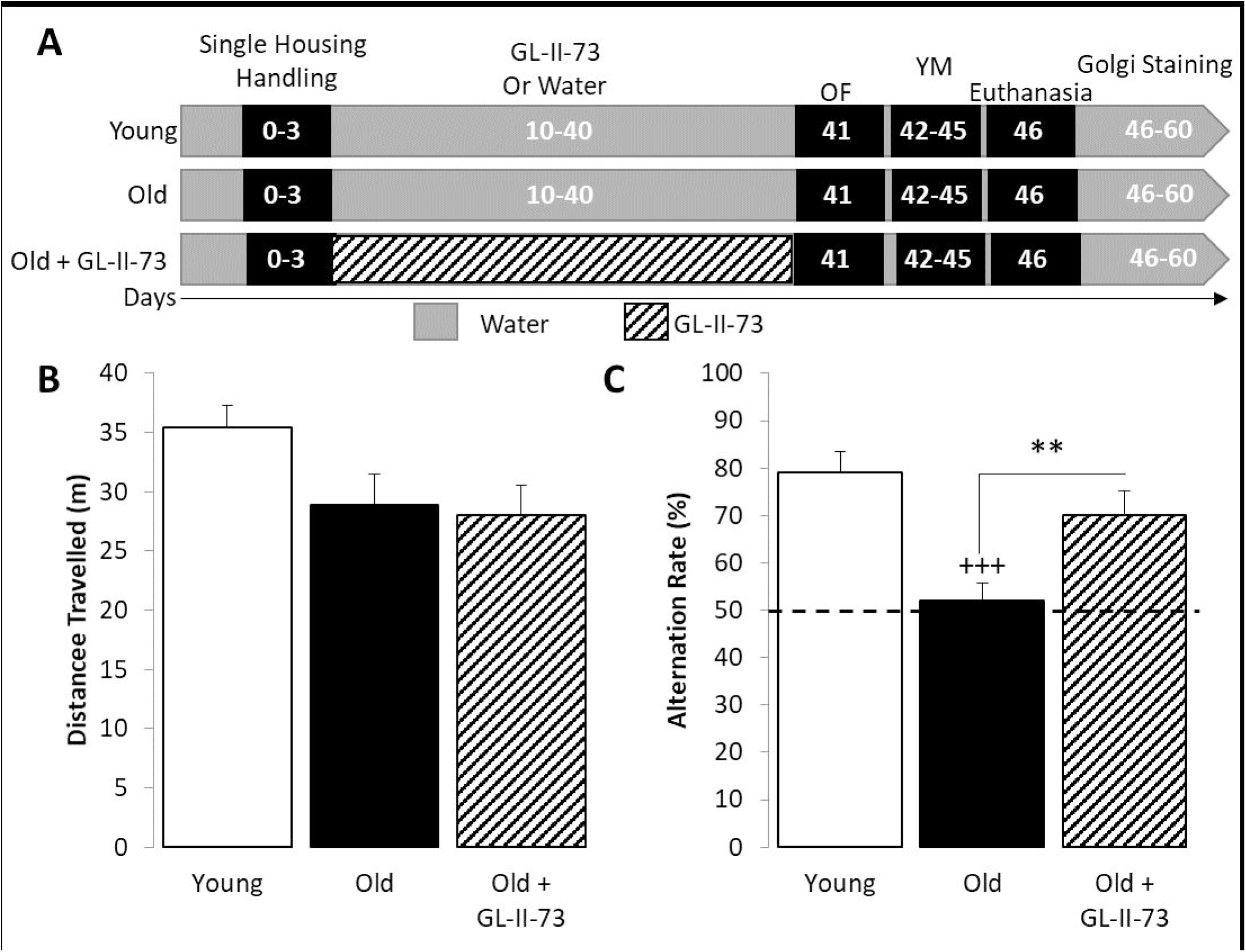
Chronic treatment with GL-II-73 reverses age-related deficit in Working memory. C57BL/6 male mice were used: young (2-month old) and old (22-month old). (**A**) Old mice received GL-II-73 in the drinking water (N=8), or water (N=14) while young mice only received water (N=11). After 30 days of treatment, mice were tested in the open-field to assess locomotor activity. (**B**) No significant differences in the total distance travelled were observed between young, old and old+GL-II-73 mice. (**C**) Mice were then trained in the alternation task in the Y Maze. ANOVA revealed a significant difference in alternation between groups, explained by a decreased alternation in old mice, compared to young, that is reversed by chronic treatment with GL-II-73. +++p<0.001 compared to “young”; **p<0.01 compared to “old”. Dash line represents chance level.

### Chronic treatment with GL-II-73 increases spatial working memory performances in old mice

After confirmation of the potential effect of GL-II-73 on dendritic morphology *in vitro*, we investigated the effect *in vivo.* Old mice received chronic administration of GL-II-73 in the drinking water for 4 weeks before being tested in the open-field to assess locomotor activity, and in the Y-Maze to assess spatial working memory (Fig.2A). ANCOVA performed on the distance travelled in the open field (Fig2.B) showed no significance differences between group (F_(2;31)_=2.4; p>0.1). ANCOVA performed on the percentage of alternation (Fig.2C) showed significant differences between groups (F_(2;25)_=11.34; p<0.001). *Posthoc* analysis identified a significant decrease of alternation with age (“Old” versus “Young”; p<0.001) that is reversed by chronic treatment of GL-II-73 (“old+GL-II-73” versus “old”, p<0.05). The “old+GL-II-73” group was not significantly different from the “young” group (p>0.1).

### Chronic treatment with GL-II-73 reverses cellular morphology changes related to aging

From each group, four mice were euthanized and brain were collected for Golgi staining. After staining, brains were shipped to NeuroDigiTech for sectioning and blinded quantification of dendritic length, spine count and spine density in the PFC and CA1 of the dorsal hippocampus. Imaging of pyramidal neurons from the PFC are presented for each group in Fig.3A., with a close-up visualization of the spines at the apical segment of the pyramidal neuron. Overall dendritic length was significantly different between group (ANOVA F_(2;143)_=5.5 p<0.05; **supplementary figure 1A**). *Post hoc* analyses showed that “old” mice had shorter dendritic length compared to “young” mice, while “old+GL-II-73” mice were neither significantly different from the “old” group nor the “young” group (p>0.05). When analyzed per dendritic segment (apical versus basal), data showed no difference between groups in the basal segment (ANOVA F_(2;71)_=1.07 p>0.05; Fig3.B), while a group difference was observed in the apical segment (ANOVA F_(2;71)_=5.3 p<0.05; Fig3.C). *Posthoc* analyses identified a significant decrease in dendritic length due to normal aging (“old” versus “young”: p<0.01) that was reversed by chronic treatment with GL-II-73 (“old+GL-II-73” versus “old”; p<0.05).

**Figure 3.**
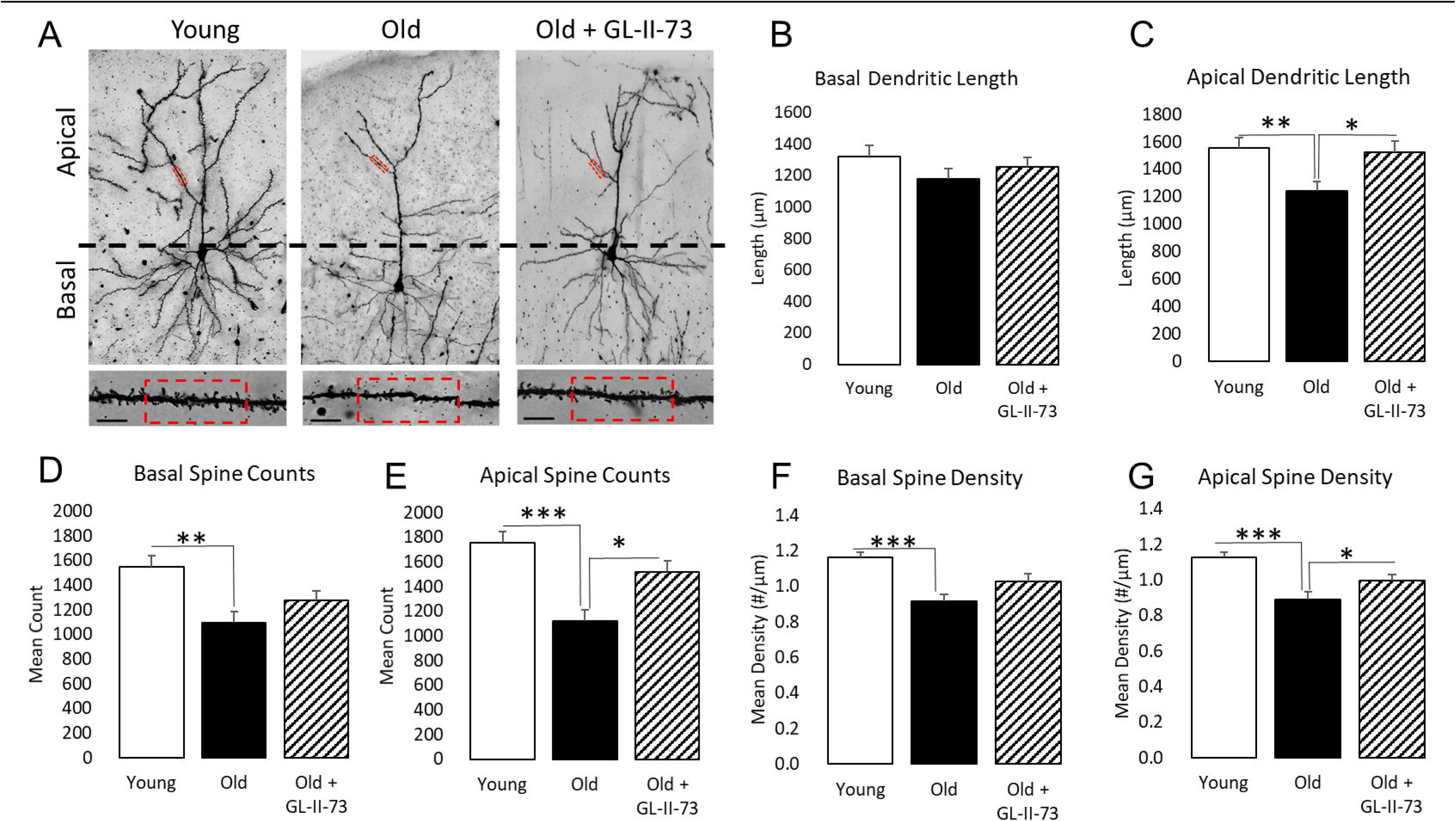
Chronic treatment with GL-II-73 reverses cellular morphology changes related to aging in the PFC. After completion of the behavioral screening, mice were euthanized and brains were stained with Golgi-Cox solution. (**A**) Pyramidal neurons (N=6) from 4 mice per group were analyzed for dendritic length, spine counts and spine density. Basal (**B**) and apical (**C**) dendritic lengths were measured. No statistical differences were observed in the basal segment, but ANOVA in the apical segment revealed significant differences between groups. This difference was explained by a decrease in dendritic length in old mice compared to young mice that was reversed by chronic treatment with GL-II-73. (**D-E**) In the same brain region, basal and apical spine counts were assessed. In the basal segment, a significant decrease was observed in old mice. In the apical segment, a similar decrease was observed with age that was reversed by chronic treatment with GL-II-73. Similarly, basal (**F**) and apical (**G**) spine density were measured. With age, a significant decrease in spine density was observed in the basal segment, and in the apical segment. However, chronic treatment with GL-II-73 only reversed this decreased in the apical segment. *p<0.05, **p<0.01, ***p<0.001 compared to “old”.

Similarly, repeated measure ANOVA performed on the total spine count showed a significant difference between groups (F_(2;143)_=16.29 p<0.001; **supplementary figure 1B**), explained by significant decrease of spine count with age (“young” versus “old”: p<0.001), reversed by chronic treatment with GL-II-73 (“old+GL-II-73” versus “old”: p<0.05). However, mice receiving the drug still had less spines than young mice (p<0.05). When looking closely at the dendritic segments, repeated measures ANOVA on the spine counts from the basal segment showed significant differences (F_(2;71)_=6.09 p<0.001, Fig3.D). *Posthoc* analyzes identified a significant decrease with age (“old” versus “young”: p<0.01) but no further significant differences. In the apical segment (Fig.3E), spine count was also significantly different between groups (repeated measure ANOVA, F_(2;71)_=10.54; p<0.001), explained by significant decrease with age (“old” versus “young”: p<0.001), reversed by chronic treatment (“old+GL-II-73” versus “old”: p<0.05). No statistical differences were observed between “young” and “old+GL-II-73” (p>0.05).

Spine density was also measured. Repeated measure ANOVA on the total spine density (**supplementary figure 1C**) showed significant differences between groups (F_(2,143)_=17.93; p<0.001), explained by decrease spine density with age (“old” versus “ young”: p<0.001), reversed by chronic treatment with GL-II-73 (“old+GL-II-73” versus “old”: p<0.01). A significant difference was observed between “old+GL-II-73” and “young” (p<0.05). When focusing on the basal segment (Fig.3F), a significant difference was identified (repeated measure ANOVA, F_(2;71)_=9.24, p<0.001) and is characterized by decreased spine density with age (“old” versus “young”: p<0.001). In the apical segment (Fig.3G), repeated measure ANOVA showed significant group differences (F_(2;71)_=10.1; p<0.001), characterized by reduced spine density due to aging, regardless of the drug treatment (“old” versus “young”, and “old+GL-II-73” versus “young”: ps<0.05).

The same analyses were carried out in the CA1 region of the dorsal hippocampus (**supplementary Figure 2**). Conversely, total dendritic length, total spine counts and spine density (**supplementary figure 3A-C**) were significantly decreased by age (repeated measure ANOVAs F_(2;143)_>9.1; ps<0.05 – *posthoc* ps<0.05), and reversed by chronic treatment with GL-II-73 (ps<0.05). Looking at dendritic length, spine counts and spine density in the basal segment (**supplementary figure 3D-F**), repeated measure ANOVAs showed significant differences (Fs_(2;71)_>3.05; ps<0.05). These differences were explained by reduced length (p<0.05), spine counts (p<0.001) and spine density (p<0.001) due to age (“old” versus “young”), and reversal of reduced spine counts (p<0.01) and spine density (p<0.05) after chronic treatment with GL-II-73.

Dendritic length, spine count and spine density in the CA1 of the hippocampus were also quantified in the apical segment of young, old and old+GL-II-73 mice (Fig.4A-C). Repeated measure ANOVAs showed significant differences between groups in all three parameters (Fs_(2;143)_>6.9; ps<0.05). *Post hoc* analyses identified these differences to be due to a significant decrease of dendritic length (p<0.01), spine counts (p<0.001) and spine density (p<0.001) with aging (“old” versus “young”). Additionally, reduced dendritic length and spine count in old mice was reversed with chronic GL-II-73 treatment (ps<0.05).

**Figure 4.**
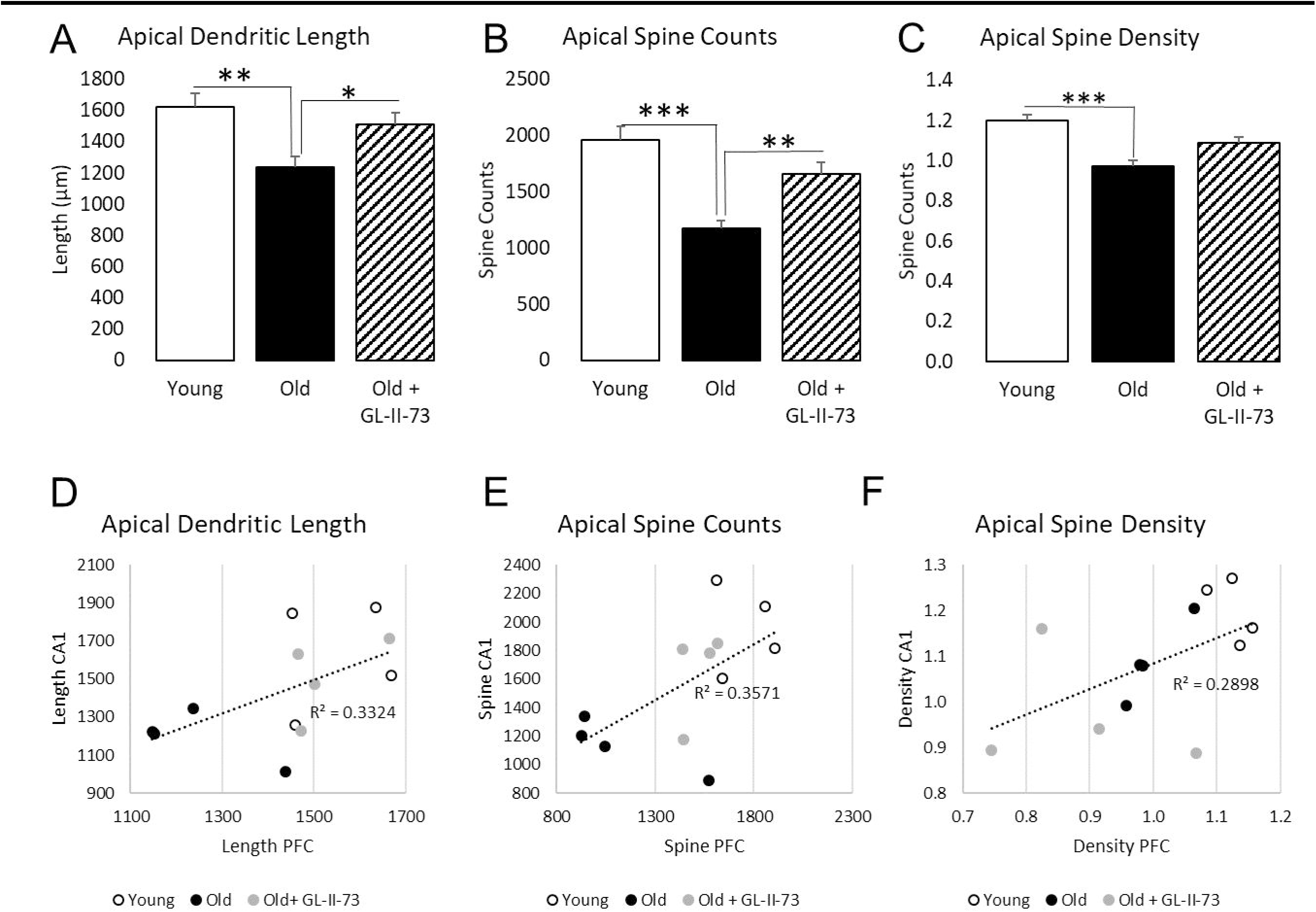
Chronic treatment with GL-II-73 reverses cellular morphology changes related to aging in the CA1, in a similar manner than the PFC. The same brains were analyzed for dendritic length, spine count and spine density in the CA1 of the hippocampus. (**A**) Dendritic length in the apical segment was significantly decreased by aging. Chronic treatment with GL-II-73 significantly reversed the reduced dendritic length in old mice. (**B**) Spine counts in the apical segment were also reduced with age. Chronic treatment with GL-II-73 reversed the reduced spine counts in old mice. (**C**) Spine density in the apical segment of pyramidal neurons in the CA1 was also significantly decreased in old mice. Chronic treatment with GL-II-73 did not significantly reversed the spine density reduction induced by aging. (**D**) Linear regression between dendritic length in the CA1 and the PFC showed positive correlation (p=0.04). (**E**) Similarly, spine count in the PFC an the CA1 were also correlated, suggesting a link between the effects observed in the PFC and the CA1. (**F**) Linear regression between the spine density in the PFC and the CA1 did not reach significance (p=0.07). *p<0.05, **p<0.01, ***p<0.001 compared to “old”.

Since the quantification of the dendritic length, spine count and spine density were performed in the PFC and CA1 of the same animals, we performed correlation analyses. Linear regression showed positive correlation between the apical dendritic lengths (Fig4.D) in the PFC and the CA1 (r=0.597; p=0.04) and between apical spine count (Fig4.E) in the PFC and CA1 (r=0.576; p=0.04). A trend was observed for the apical spine density (Fig.4F; r=0.5383; p=0.07).

### Cessation of treatment alters the pro-cognitive efficacy, but does not reduce the cellular effects

A separate cohort of mice was tested in the Y maze after 30 days of treatment with GL-II-73, or after 30 days of treatment and a subsequent washout period of 1 week (“washout” group; Fig5.A). ANCOVA performed on the alternation rate showed significant differences between groups (F_(3;17)_=9.48; p=0.007), explained by significant decrease of alternation with aging (“old” versus “young”: p<0.01), reversed by chronic treatment (“old+GL-II-73” versus “old”:p<0.01). However, mice in the “washout” group had decreased alternation rate compared to “young” and “old+GL-II-73” (ps<0.01; Fig.5B).

**Figure 5.**
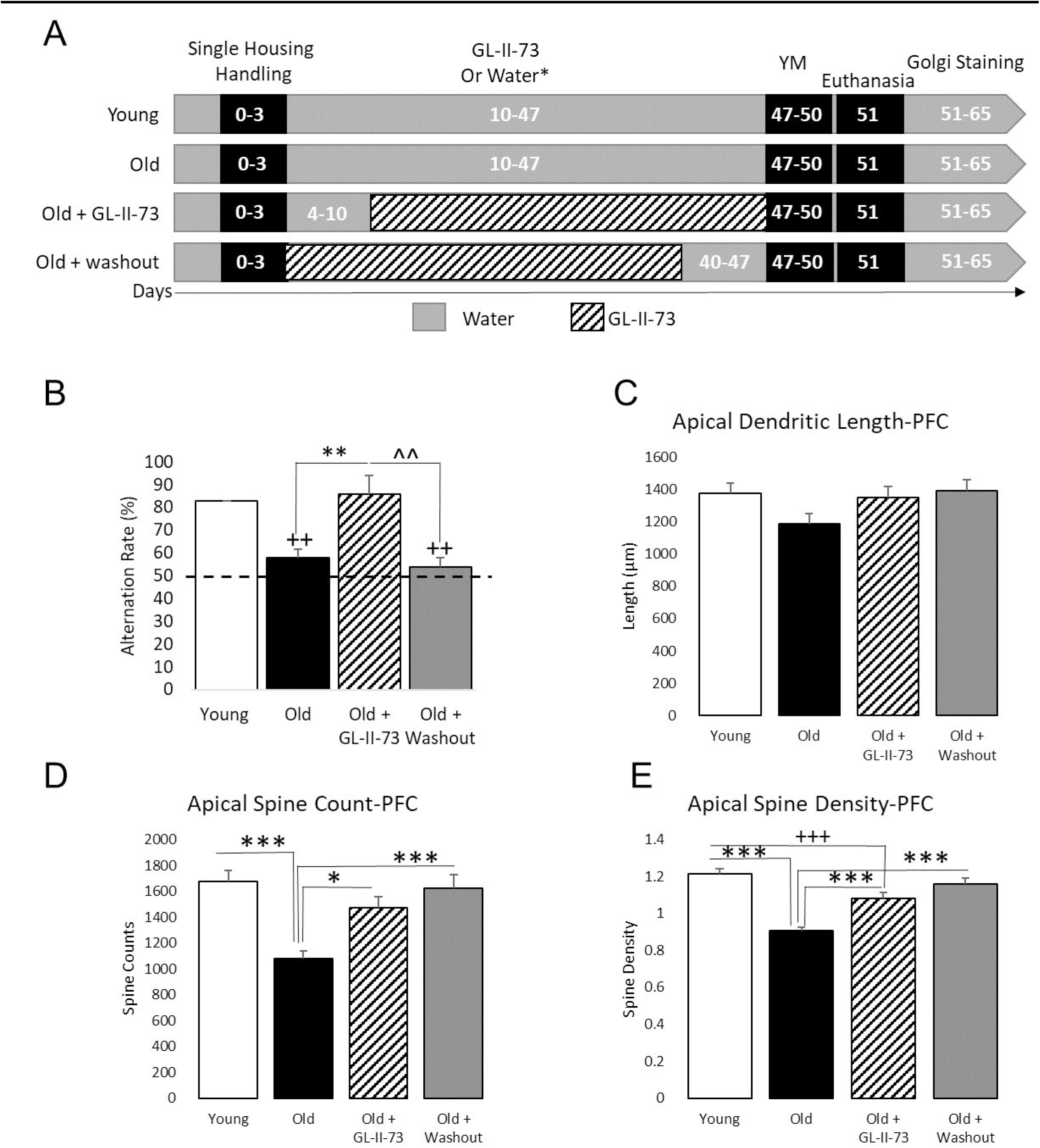
Cessation of chronic treatment with GL-II-73 limits the precognitive efficacy but does not alter the effect on dendritic morphology. C57BL/6 male mice were used: young (2-month old) and old (22-month old). (**A**) Old mice received GL-II-73 in the drinking water (N=6), or water (N=6), or GL-II-73 for 30 days followed by 7 days of washout (N=6) while young mice only received water (N=8). (**B**) After completion of the treatment schedule, mice were trained in the alternation task in the Y Maze. ANOVA revealed a significant difference in alternation between groups, explained by a decreased alternation in old mice, compared to young, that is reversed by chronic treatment with GL-II-73. However, the 7-day washout limited the precognitive effect. After completion of the behavioral testing, mice were euthanized, brains were collected and stained for morphology analyzes, and apical morphology features were measured. (**C**) ANOVA performed on the apical dendritic length did not reach significance (p>0.05). (**D**) ANOVA performed on the apical spine count reached significance (p<0.05), and could be explained by a decrease of spine count with age, that is reversed with chronic treatment with GL-II-73, even after a washout period. (**E**) Similar findings were obtained with apical spine density measurement, where age decreased spine density, but chronic treatment with GL-II-73 reversed this deficit, even after a 7-day washout. ++p<0.01, +++p<0.001 compared to “young”; *p<0.05, **p<0.01, ***p<0.001 compared to “old”; ˄˄p<0.01 compared to “old+GL-II-73”. Dash line represents chance level.

After completion of the behavioral testing, mice were euthanized and brains collected for Golgi staining (**supplementary Figure 4**). ANOVA performed on total dendritic length, spine counts and spine density in the PFC were performed (Fs_(3,188)_>3.5; ps<0.05; **Supplementary Figure 5**). These differences were explained by a decrease of dendritic length in old mice, compared to young mice (p<0.05). Regarding the total spine counts, old mice showed decreased spine counts compared to young (p<0.001), that was reversed by chronic treatment with GL-II-73 (“old+GL-II-73” versus “old”; p<0.001) even after a 1-week washout (“old+washout” versus “old”; p<0.001). The significant differences in the total spine density were explained by reduced spine density in old mice compared to young (p<0.001), partially reversed by chronic treatment, even with a 1-week washout (“old” versus “old+GL-II-73”, “old” versus “old+washout”, “young” versus “old+GL-II-73” and “young” versus “ old+washout”: ps<0.05). In the apical segment, ANOVA on the total dendritic length did not reach significance (p=0.1; Fig.5C). ANOVAs performed on the apical spine counts and spine density reached significance (F_(3,188)_>9.6; p<0.001). The significant difference in the apical spine count (Fig.5D) was explained by reduced spine count during aging (“old” versus “young”: p<0.001), reversed by chronic treatment with GL-II-73 (“Old+GL-II-73” versus “old”:p<0.05), even with a 1-week washout (“old+washout” versus “old”: p<0.001). Finally, the significant difference in apical spine density (Fig.5E) in the PFC was explained by a significant decrease with age (“old” versus “young”:p<0.001), partially reversed by chronic treatment with GL-II-73 (“old+GL-II-73” versus “old” and “young”; ps<0.01), and fully reversed by chronic treatment with a 1-week washout (“old+washout” versus “old”:p<0.001). In the basal segment, ANOVAs performed on the spine counts and the spine density showed significant differences (F_(3,188)_>9.1; ps<0.001) but not on the dendritic length (F_(3,188)_>2.3; p=0.07) (**supplementary Figure 5D-F**). Spine count and density in the basal segment were decreased with aging (“old” versus “young”: ps<0.001), and reversed by chronic treatment (“old+GL-II-73” versus “old”: ps<0.05), even with a 1-week washout (“old+washout” versus “old”: ps<0.05).

Similar analyses were performed on the dendritic length, spine count and spine density in the CA1 of the same animals (**supplementary Figure 6**). The data obtained in the CA1 confirmed what was shown in the PFC. There were no significant differences in total, basal or apical dendritic length (data not shown). A significant difference was found in total spine count (F_(3,188)_=8.95; p<0.001), explained by a decrease in spine count with age (“old” versus “young”: p<0.001), that was significantly reversed only in the washout group (“old+washout” versus “old”: p<0.001; **Supplementary Figure 7A**). The same effects were observed when looking specifically at the basal and apical segment (**Supplementary Figure 7B-C**). Total spine density was also showing significant differences (ANOVA F_(3,188)_=25.65, p<0.001; **supplementary Figure 7D**), explained by decreased density with aging (“old” versus “young”: p<0.001), partially reversed by chronic treatment (“Old+GL-II-73” versus “old” and “young”: ps<0.01), even with a 1-week washout (“old+washout” versus “old” and “young”: ps<0.05). The decrease in spine density with age was confirmed in the basal and apical segments (ANOVAs Fs_(3,188)_>10.5; ps<0.01), as well as the reversal with chronic treatment with GL-II-73, even with a 1-week washout (**Supplementary Figure 7E-F**).

## Discussion

This study was based on the observation that normal aging induces neuronal loss contributing to cognitive decline. Here, we first confirmed that normal aging in mice induces cognitive deficits, with old mice exhibiting impaired working memory in the alternation task, consistent with previous finding (59, 63, 64). Many compounds (65), drugs (66), natural extracts (64) or physical exercises (67) demonstrated potential at reversing cognitive decline in animal models, but with little-to-no success in human populations (68, 69). This lack of efficacy can partially be explained by the focus of such approaches aiming at reversing the symptoms more than the underlying pathology. Indeed, it is unclear at the moment if treatment avenues that show pro-cognitive efficacy actually alleviate the underlying pathology, such as neuronal dysfunction or neuronal shrinkage. Companion tools, like radio-ligands for PET imaging can be developed to demonstrate such efficacy, but priority is still given at alleviating the symptoms.

In the context of aging, the “morphomolecular” senescence has a broad impact on different systems, including the GABAergic system. Reduced GABAergic functions are a hallmark of aging, and previous studies from our group demonstrated that targeting α_5_-GABAA receptors to bypass such deficits could reverse age-related cognitive decline (59). In the present study, we investigated the potential combined efficacy of GL-II-73 on symptoms and underlying pathology, i.e. neuronal shrinkage. We demonstrated that chronic treatment with GL-II-73 not only reverses cognitive decline, but also reverses dendritic shrinkage and spine atrophy in the PFC and the CA1. Indeed, chronic treatment with GL-II-73 increased dendritic length, spine count and spine density in the both brain regions, in old mice. To our knowledge, it is the first time that a drug acting on the GABAergic system demonstrates neurotrophic effects, while commonly used benzodiazepines seem to decrease spine density in cortical pyramidal neurons (70). Other drugs have shown neurotrophic effects (71), such as ketamine for the treatment of depressive episodes (72). Recent studies have shown that the antidepressant action of ketamine is mediated by its activity on NMDA receptor of GABA neurons (73, 74). Further studies will be needed to identify what could be the mechanism of action of GL-II-73 allowing this neurotrophic effect, and if it has shared pathways with ketamine.

Interestingly, the neurotrophic effect of GL-II-73 was more pronounced in the apical segment of the pyramidal neuron dendrites compared to the basal segment. It is known that the α_5_-GABA-A receptors are located in the apical segment of the pyramidal neurons (52), so these results suggest that the α_5_-GABA-A receptors directly mediate this neurotrophic effect. However, a significant increase of spine counts and spine density in basal segment of pyramidal neurons of the CA1 from the treatment group was also observed. This effect can either suggest the potential presence of the α_5_-GABA-A receptors in the basal segment of pyramidal neurons of the CA1, or a generalized effect happening downstream of the apical effect. Such hypotheses would need to be validated but is consistent with previous finding showing that the α_5_-GABA-A receptors are involved in the dendritic outgrowth and spine maturation of pyramidal neurons (56).

When the drug is no longer given to the animals, the neurotrophic effect remains but the pro-cognitive effect diminishes. One could argue that the drug needs to be on board to improve cognitive performances continuously, even in the event of a strong neurotrophic effect. This would suggest that the neurotrophic effect alone is not sufficient to recover normal cognitive processes, and suggests that the “rejuvenated” neurons are not as functional as neurons of younger animals. Here, we did not investigate if the “rejuvenated” neurons were similarly functional compared to neurons from younger animals, or if their functions were more similar to neurons from old animals. Using slice electrophysiology could answer thia question and investigate if these neurons were truly rejuvenated or if the neurotrophic effect was limited to morphological features. In addition, the present study only tested 1 treatment duration (4 weeks) and 1 washout period (1 week). Investigating if longer treatment with GL-II-73 improves the stability of the cognitive functions when the treatment is spotted could be a solution to improve the functions of the rejuvenated neurons.

Despite promising results in mice suggesting a relief of the cognitive symptoms and a rejuvenation of neurons in brain regions involved in cognitive processes, it would be naïve to think that a single drug could reverse all the changes induced by normal aging. However, GL-II-73 is the first drug targeting an inhibitory system that shows neurotrophic, associated with symptom relief. With its potential anxiolytic and antidepressant effects (59), GL-II-73 could become a game changer for patients suffering from cognitive decline due to GABAergic downregulation. In the context of aging, and early stages of Alzheimer’s disease, GL-II-73 could possibly be paired with Aducanumab(35), a potent monoclonal antibody directed against β-amyloid oligomers, to further slowdown the progression of the disease and strengthen neuronal “health” and cognitive functions. Further testing in other cognitive domains, and other models (Alzheimer’s disease, neurodegenerative disease etc.) will demonstrate the validity of this new drug for future therapeutic uses.

