## Supplementary for "Chronic administration of a positive allosteric modulator at the α5-GABAA receptor reverses age-related dendritic shrinkage"

### --- SUPPLEMENTARY FIGURES ---

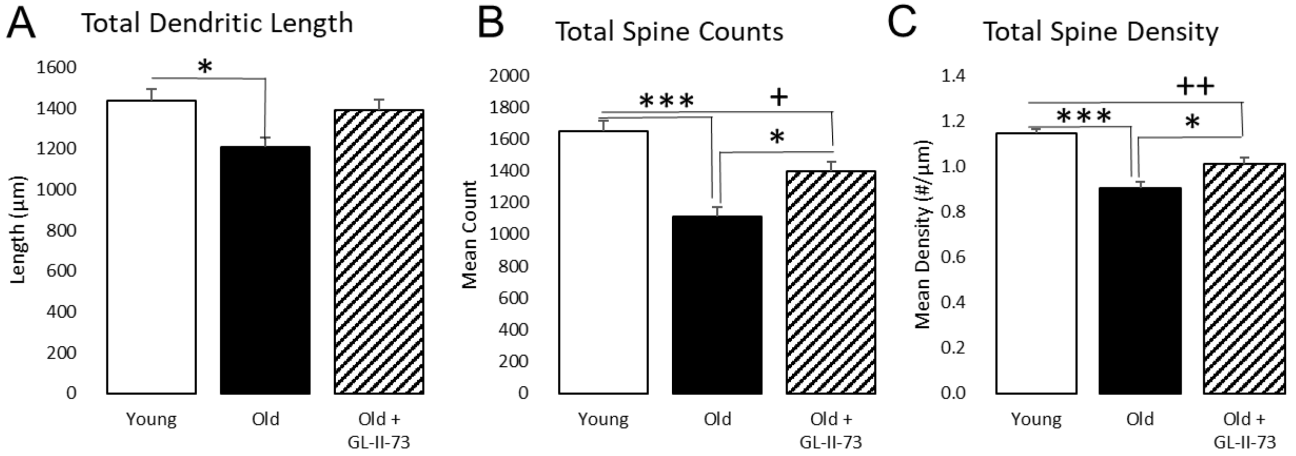


**Supplementary Figure 1. Total dendritic length, total spine counts and total spine density in the PFC of young, old and old+GL-II-73 mice.**

(**A**) A significant decrease in dendritic length was observed in old compared to young. (**B**) A significant decrease in total spine number was observed in old mice compared to young, and was partially reversed by chronic treatment with GL-II-73. (**C**) Similar findings were obtained with the total spine density in the PFC. Old mice showed decreased spine density compared to young mice. This deficit was partially reversed by chronic treatment with GL-II-73. +p<0.05, ++p<0.01 compared to "young"; *p<0.05, ***p<0.001 compared to "old".

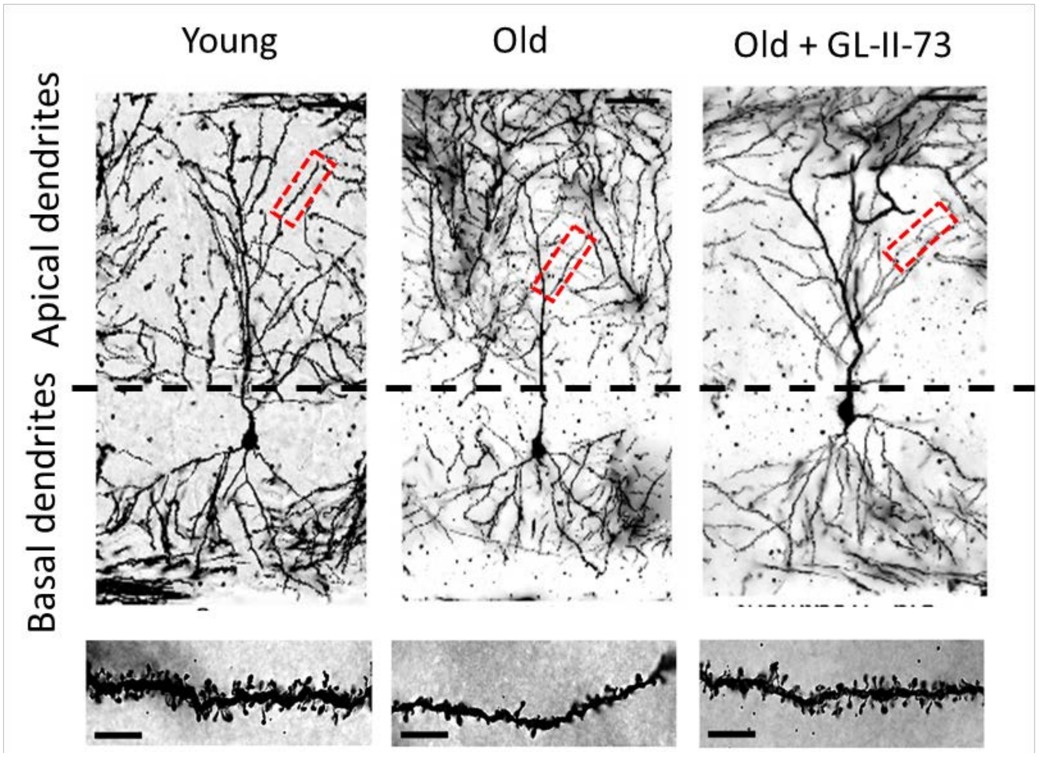


**Supplementary Figure 2: Representative pictures of pyramidal neurons in the CA1 of the hippocampus of mice, depending on the treatment group.**

Brains from young, old and old treated mice were harvested and stained using a Golgi-Cox technique. Sections were mounted on slides and imaged at 100X for 3D dendritic reconstruction, followed by counting of the spines throughout the entire dendritic trees. Scale bar represents 50µm.

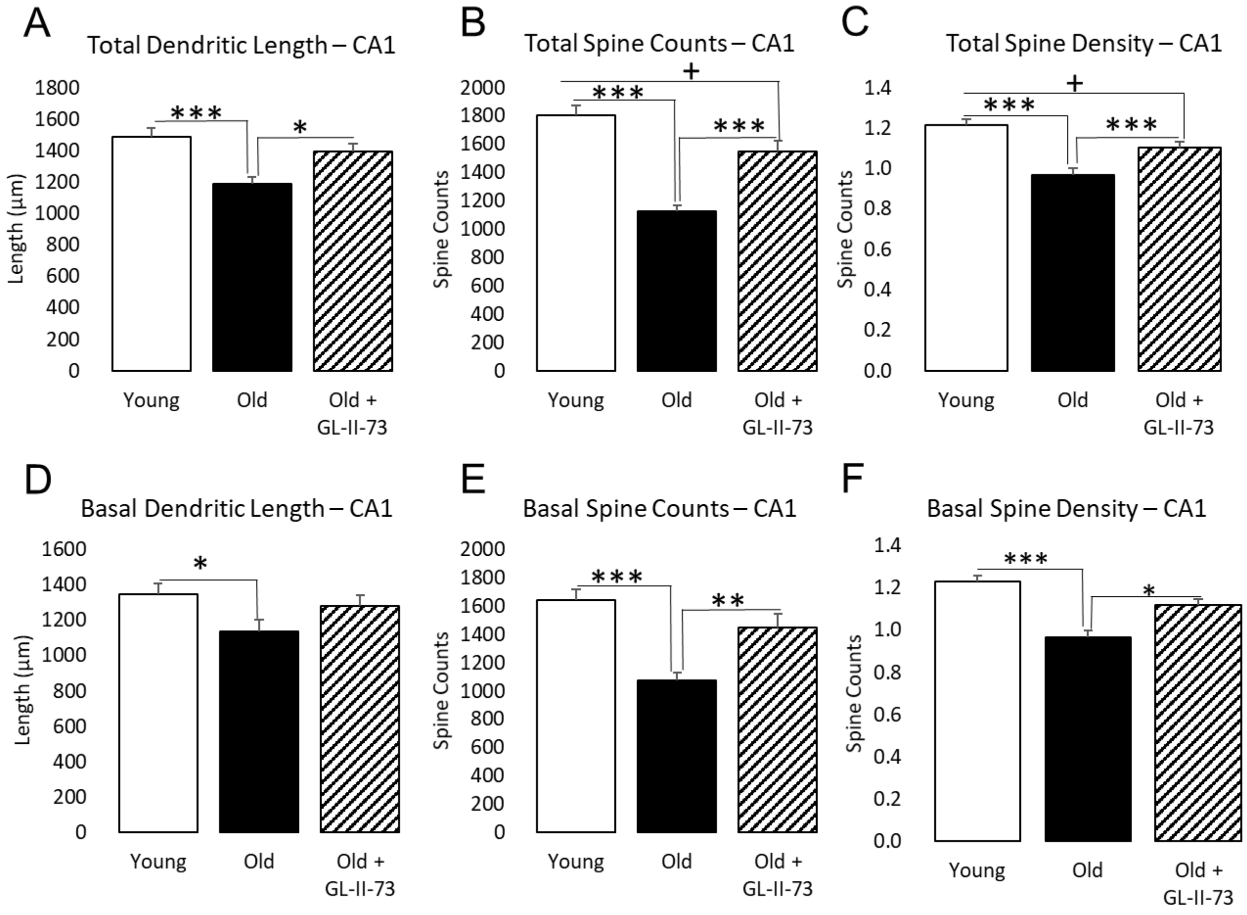


**Supplementary Figure 3: Morphological features of the pyramidal neurons of the CA1 of the hippocampus depending on the treatment group**

(**A**) Total dendritic length of pyramidal neurons in the CA1 showed a significant decrease with age, that is reversed by chronic treatment with GL-II-73. (**B**) Total spine count was also significantly decrease in the CA1 of old mice, and such decrease was partially reversed by chronic treatment with GL-II-73. (**C**) Similar findings were obtained regarding the total spine density, with a significant decrease due to age, that is partially reversed by chronic treatment with GL-II-73. (**D**) With a focus on the basal segment, total dendritic length was decrease with age. Basal dendritic length of the “old+GL-II-73” group was not significantly different from the “old” or “young” groups. (**E**) Basal spine counts were decreased with age, and reversed with chronic treatment with GL-II-73, as well as (**F**) basal spine density. +p<0.05 compared to “young”; *p<0.05, **p<0.01, ***p<0.001 compared to “old”.

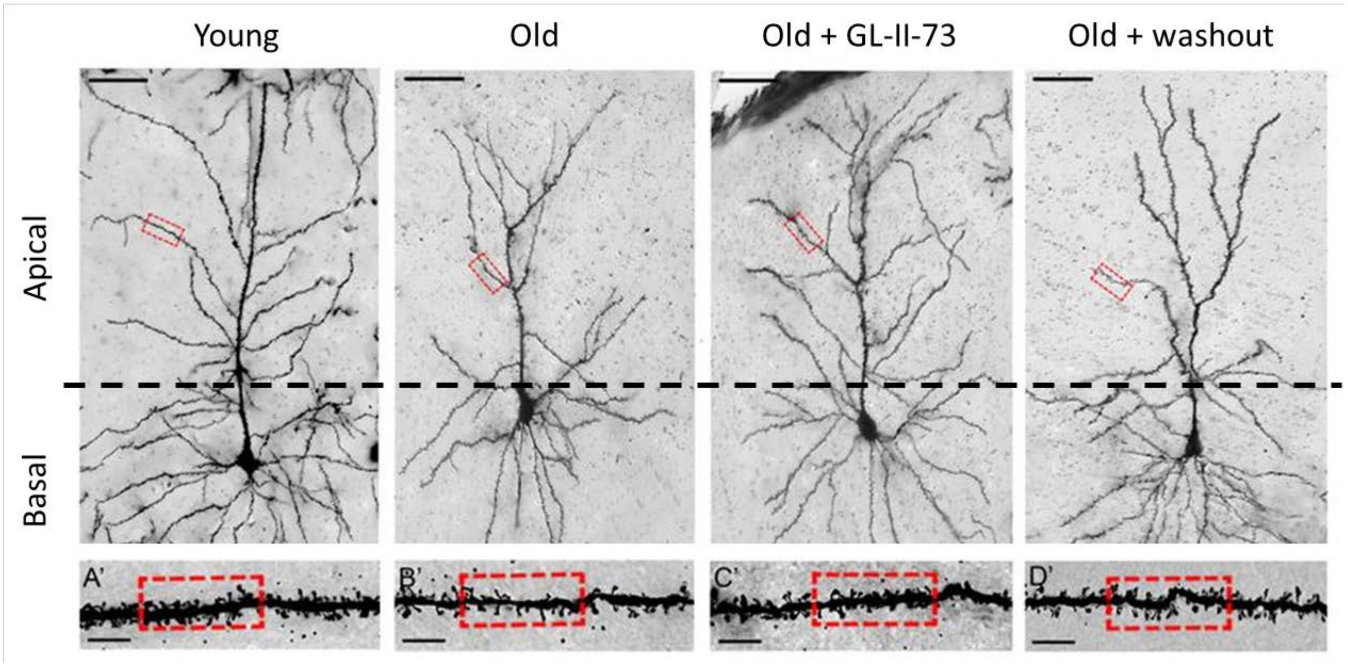


**Supplementary Figure 4: Representative pictures of pyramidal neurons in the PFC of mice, depending on the treatment group.**

Brains from young, old, old+GL-II-73 and old+washout mice were harvested and stained using a Golgi-Cox technique. Sections were mounted on slides and imaged at 100X for 3D dendritic reconstruction, followed by counting of the spines throughout the entire dendritic trees. Scale bar represents 50µm.

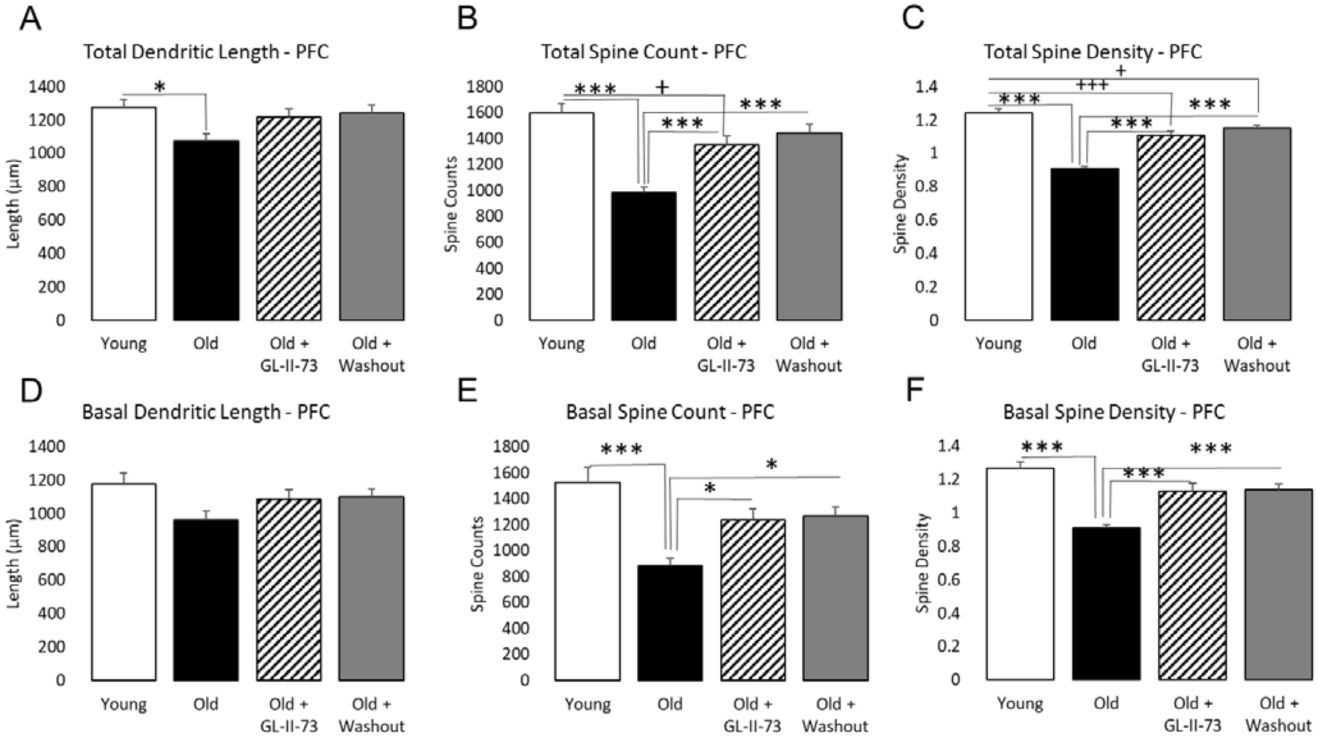


**Supplementary Figure 5: Morphological features of the pyramidal neurons of the PFC depending on the treatment group**

(**A**) Total dendritic length of pyramidal neurons in the PFC showed a significant decrease with age. Both "old+GL-II-73" and "old+washout" group were not different from the "old" and "young" groups. (**B**) Total spine count was also significantly decrease in the PFC of old mice, and such decrease was partially reversed by chronic treatment with GL-II-73, and totally reversed with chronic treatment with GL-II-73 and a 7-days washout. (**C**) Similar findings were obtained regarding the total spine density, with a significant decrease due to age that is partially reversed by chronic treatment with GL-II-73, with or without a 7-days washout. (**D**) With a focus on the basal segment, total dendritic length was not significantly affected. (**E**) Basal spine counts were decreased with age and reversed with chronic treatment with GL-II-73, with or without a 7-days washout. (**F**) Similar conclusion also applies to the basal spine density measurement, with a significant decrease with age that is reversed by chronic treatment with GL-II-73, regardless of the washout period. +$p<0.05$, +++$p<0.001$ compared to "young"; *$p<0.05$, **$p<0.01$, ***$p<0.001$ compared to "old".

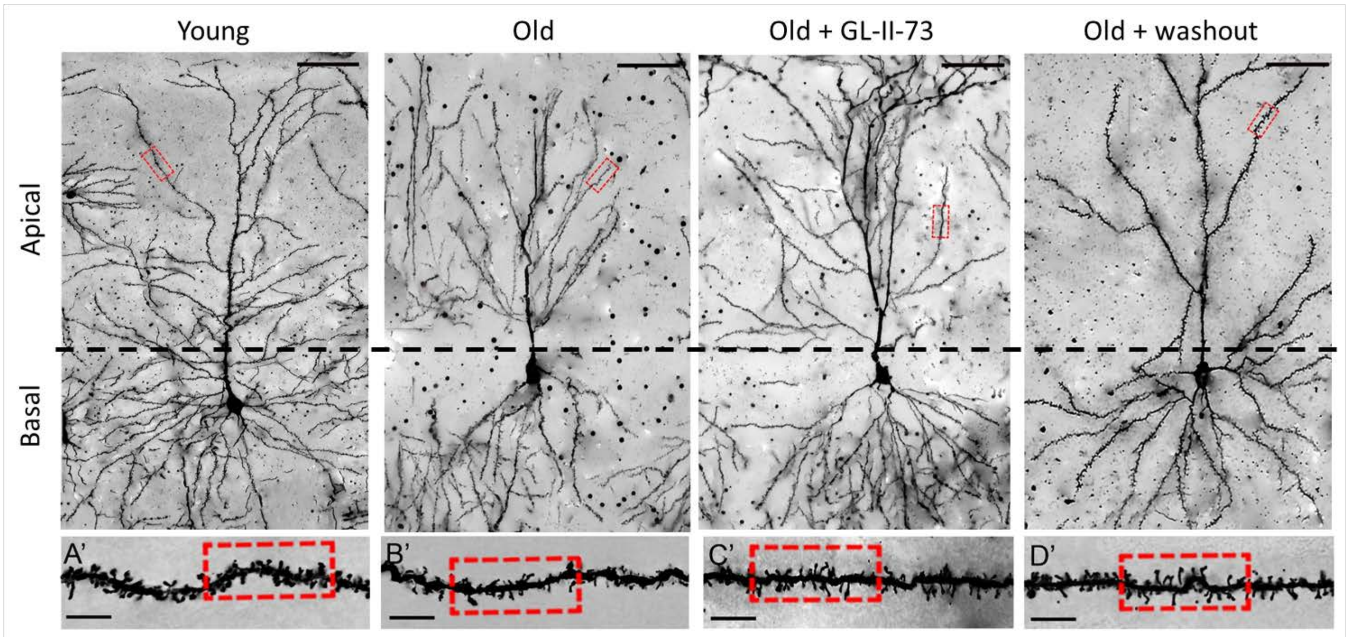


**Supplementary Figure 6: Representative pictures of pyramidal neurons in the CA1 of the hippocampus of mice, depending on the treatment group.**

Brains from young, old, old+GL-II-73 and old+washout mice were harvested and stained using a Golgi-Cox technique. Sections were mounted on slides and imaged at 100X for 3D dendritic reconstruction, followed by counting of the spines throughout the entire dendritic trees. Scale bar represents 50µm.

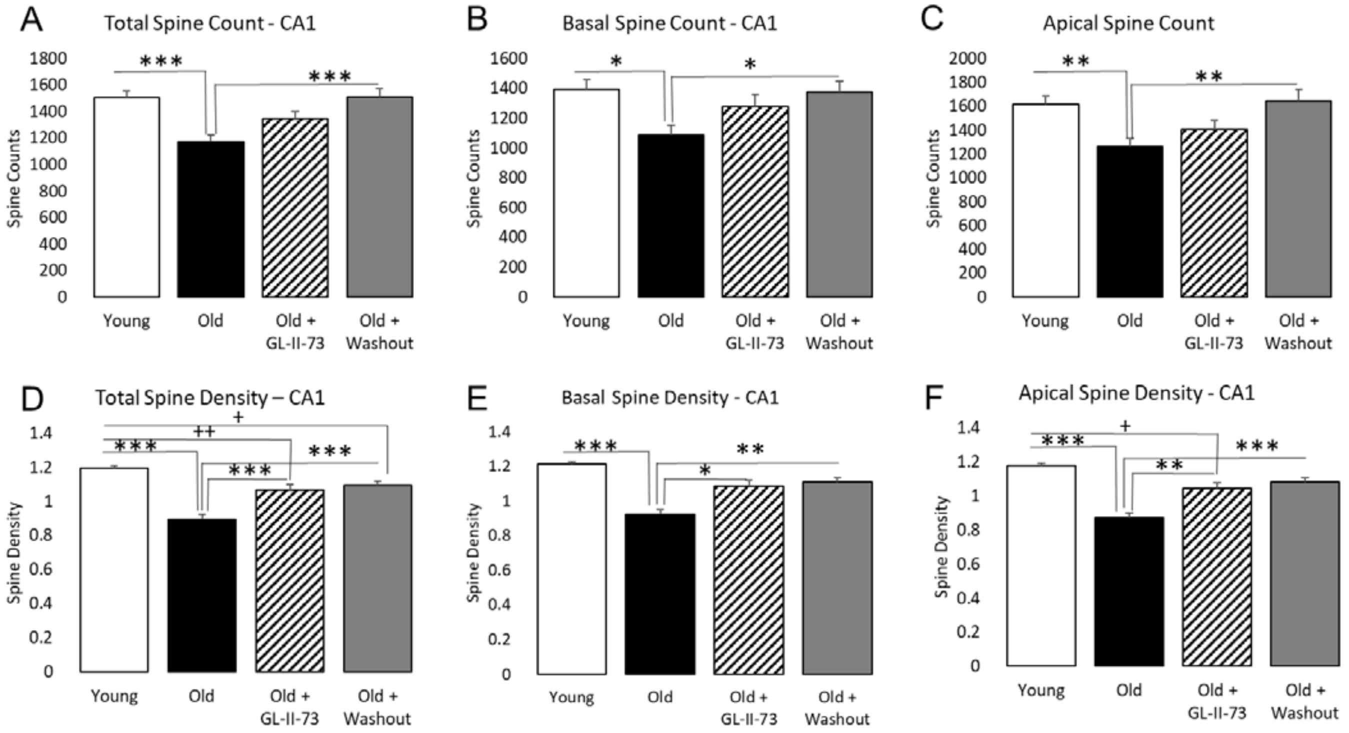


**Supplementary Figure 7: Morphological features of the pyramidal neurons of the CA1 of the hippocampus depending on the treatment group**

(**A**) Total spine counts of pyramidal neurons in the CA1 showed a significant decrease with age, significantly reversed only in the "old+washout" group. The "old+GL-II-73" group was not different from the "old" and "young" groups. Basal (**B**) and apical spine count (**C**) showed similar profiles in the CA1. (**D**) Total spine density was significantly decreased in the CA1 of old mice, and such deficit was partially reversed by chronic treatment with GL-II-73, with or without a 7-days washout. (**E**) Basal spine density was decreased with age and reversed with chronic treatment with GL-II-73, with or without a 7-days washout. (**F**) Apical spine density was decreased with age, partially reversed with GL-II-73 treatment, and totally reversed in the "washout" group. +$p<0.05$, ++$p<0.01$ compared to "young"; *$p<0.05$, **$p<0.01$, ***$p<0.001$ compared to "old".
